# Persistent lung inflammation and alveolar-bronchiolization due to Notch signaling dysregulation in SARS-CoV-2 infected hamster

**DOI:** 10.1101/2024.05.13.593878

**Authors:** Can Li, Na Xiao, Wenchen Song, Alvin Hiu-Chung Lam, Feifei Liu, Xinrui Cui, Zhanhong Ye, Yanxia Chen, Peidi Ren, Jianpiao Cai, Andrew Chak-Yiu Lee, Honglin Chen, Zhihua Ou, Jasper Fuk-Woo Chan, Kwok-Yung Yuen, Hin Chu, Anna Jin-Xia Zhang

**Affiliations:** State Key Laboratory of Emerging Infectious Diseases, Carol Yu Centre for Infection, Department of Microbiology, Li Ka Shing Faculty of Medicine, The University of Hong Kong, Pokfulam, Hong Kong Special Administrative Region, China; Centre for Virology, Vaccinology and Therapeutics, Hong Kong Science and Technology Park, Hong Kong Special Administrative Region, China; Department of Clinical Microbiology and Infection Control, The University of Hong Kong-Shenzhen Hospital, Shenzhen, China; Department of Microbiology, Queen Mary Hospital, Pokfulam, Hong Kong Special Administrative Region, China; Shenzhenkey Laboratory of Unknown Pathogen Identification, BGI Research, Shenzhen, China

**Author notes:** **Correspondence:** Hin Chu, Department of Microbiology, Li Ka Shing Faculty of Medicine, The University of Hong Kong, Pokfulam, Hong Kong Special Administrative Region, China; Anna JX Zhang, Department of Microbiology, Li Ka Shing Faculty of Medicine, The University of Hong Kong, Pokfulam, Hong Kong Special Administrative Region, China. These authors are co-first authors (Can Li, Na Xiao, Wenchen Song, Alvin Hiu-Chung Lam, Feifei Liu).

**Keywords:** hamster, PASC, long COVID-19, SARS-CoV-2, basal cell, alveolar-bronchiolization, Notch

## Abstract

Long COVID or Post-acute sequalae of COVID-19 (PASC) defines the persistent signs, symptoms, and conditions long after initial SARS-CoV-2 infection which affecting over 10% of COVID-19 patients, with 40% of them affecting respiratory system. The lung histopathological changes and underlying mechanism remain elusive. Here we systemically investigate histopathological and transcriptional changes at 7, 14, 42, 84 and 120 days-post-SARS-CoV-2-infection (dpi) in hamster. We demonstrate persistent viral residues, chronic inflammatory and fibrotic changes from 42dpi to 120dpi. The most prominent lung histopathological lesion is multifocal alveolar-bronchiolization observed in every animal from 14dpi until 120dpi. However, none of the above are observed in hamsters recovered from influenza A infection. We show airway progenitor CK14+ basal cells actively proliferate, differentiate into SCGB1A+ club cell or Tubulin+ ciliated cells, leading to alveolar-bronchiolization. Most importantly, Notch pathway is persistently upregulated. Intensive Notch3 and Hes1 protein expression are detected in alveolar-bronchiolization foci, suggesting the association of sustained Notch signaling with dysregulated lung regeneration. Lung spatial transcriptomics show upregulation of genes positively regulating Notch signaling is spatially overlapping with alveolar-bronchiolization region. To be noted, significant upregulation of tumor-related genes was detected in abnormal bronchiolization region by spatial transcriptomics analysis, indicating possible risk of lung carcinoma. Collectively, our data suggests SARS-CoV-2 infection caused chronic inflammatory and fibrotic tissue damages in hamster lung, sustained upregulation of Notch pathway signaling contributed to the dysregulated lung regeneration and CK14+ basal cells-driven alveolar-bronchiolization. The study provides important information for potential therapeutic approaches and probable long-term surveillance of malignancy in PASC management.

## Introduction

Severe acute respiratory syndrome coronavirus 2 (SARS-CoV-2) caused more than 775 million infections and over 7.04 million deaths (*1*). Many patients recovered from SARS-CoV-2 infection experience symptoms of long COVID or post-acute sequalae of COVID-19 (PASC) (*2*), which include dyspnea, chest pain, palpitations, nausea, abdominal pain, headache, sub-fever, fatigue, depression, anxiety and cognitive dysfunction (*2, 3*). Among these, respiratory symptoms are reported in 5-40% of patients with PASC (*2, 3*). Dyspnea and persistent cough were found in up to 40% and 20% of patients after 4-8 months (*2, 3*). Recent study identified 79.4% of hospital-discharged patients with more than 10% involvement of residual lung abnormalities after 8 months (*4*). CT scans showed fibrosis features in lung, including ground-glass opacification and reticulation (*4–6*).

The pulmonary abnormalities and underlying mechanisms in PASC are still not fully understood. Current hypothesis of mechanisms causing chronic lung damages includes viral persistence, fibrosis, and dysregulated lung regeneration (*7, 8*). Viral persistence induces inflammation and possible fibrosis. Incomplete or prolonged alveoli and airway regeneration will lead to impaired capacity of gas exchange. It’s crucial to elucidate lung regeneration process in PASC. SARS-CoV-2 infection causes diffuse alveolar damage (DAD) with massive alveolar type (AT)1 and AT2 cell death (*9*), which activate multiple pulmonary progenitor cells for tissue repair along the resolution of acute inflammation (*10*). Upon mild injury/infection, AT2 cells are stimulated to multiply for self-renewal and trans-differentiate into AT1 cells (*11*). Upon severe injury, multiple regional lung progenitor cells contribute to the lung repair, including basal cells, club cells, bronchioalveolar stem cells (BASCs) and AT2 cells (*9–13*). Large number of studies suggest basal cells play important role in lung regeneration upon severe injury, e.g. bleomycin/LPS injury and influenza virus infection (*14–17*). In airways, basal cells respond rapidly after ciliary loss. Basal cells survived and differentiated into ciliated cells in SARS-CoV-2 infected bronchial organoids (*18*). CK5+ basal cells also actively proliferated and involved in alveolar repair in mice and human after influenza A and SARS-CoV-2 infection (*12, 19*). On the other hand, aberrant basal cells activities may lead to dysregulated differentiation, which is associated with chronic obstructive pulmonary diseases (COPD) and pre-neoplastic lesions (*20, 21*). However, the role of basal cells in alveolar regeneration and lung abnormalities after SARS2-CoV-2 infection have not been studied in detail.

Serial clinical lung tissue samples are not available to provide comprehensive information in PASC patients. We previously established and demonstrated that Syrian golden hamster is a physiological relevant small animal model for SARS-CoV-2 infection (*22*). Moreover, it is superior to K-18 mouse and ferret for study of the long-term sequelae with lung recovery after SARS-CoV-2 induced non-lethal disease in hamster model (*22–24*). Here through studying serial histopathological changes in hamsters after SARS-CoV-2 infection from 7dpi to 120dpi, we demonstrate persistent viral residues, chronic inflammation, fibrosis and dysregulated lung regeneration. We further prove that CK14+ basal cells in distal airways serves as major progenitors of bronchioles epithelium regeneration, while CK14+ basal cells activation prolongs, with proliferation in alveoli and differentiation to club cells and ciliated cells, leading to multiple abnormal bronchiolization foci in the lung. The possible mechanism for CK14+ cell-driven alveoli regeneration might be mediated by continuous high expression of Notch signaling pathway. Collectively, this study provides evidence of viral persistence, chronic inflammatory and fibrotic changes in the lung together with persistent alveolar-bronchiolization, which may contribute to the long-lasting respiratory symptoms in human. Moreover, our study addresses the potential carcinogenesis in lung after SARS-CoV-2 infection and calls for further investigations and long-term surveillance of malignancy.

## Results

### Prominent chronic histopathological damages in hamster lung after acute SARS-CoV-2 infection

To characterize the temporal lung histopathology after SARS-CoV-2 infection, we conducted a longitudinal study in 6-8 weeks old hamsters challenged with 10^3^ PFU of SARS-CoV-2 wild-type strain HK-13 till 120dpi (Fig 1a). The most prominent histological feature of the lung at 120dpi was abundant foci of alveolar hyperplasia (Fig 1b). Closer examination showed alveolar consolidation, increased thickness of alveolar septa with reduced air sac, immune cell infiltration in peribronchiolar and perivascular connective tissues (Fig 1b). Collapse of alveoli was obvious. Pleural fibrosis was also observed in large proportions of the infected hamsters (Fig 1b). Multiple alveolar hyperplasia foci distributed from around bronchioles to distal lung, affecting 4 to 11.3% of lung examined. The cellular morphologies in these abnormal foci were mainly cuboidal or columnar shape, which is consistent with the characteristics of alveolar-bronchiolization (Fig 1b). None of these changes were observed in age-matched mock control hamsters, which strongly indicated their association with previous SARS-CoV-2 infection.

**Figure 1.**
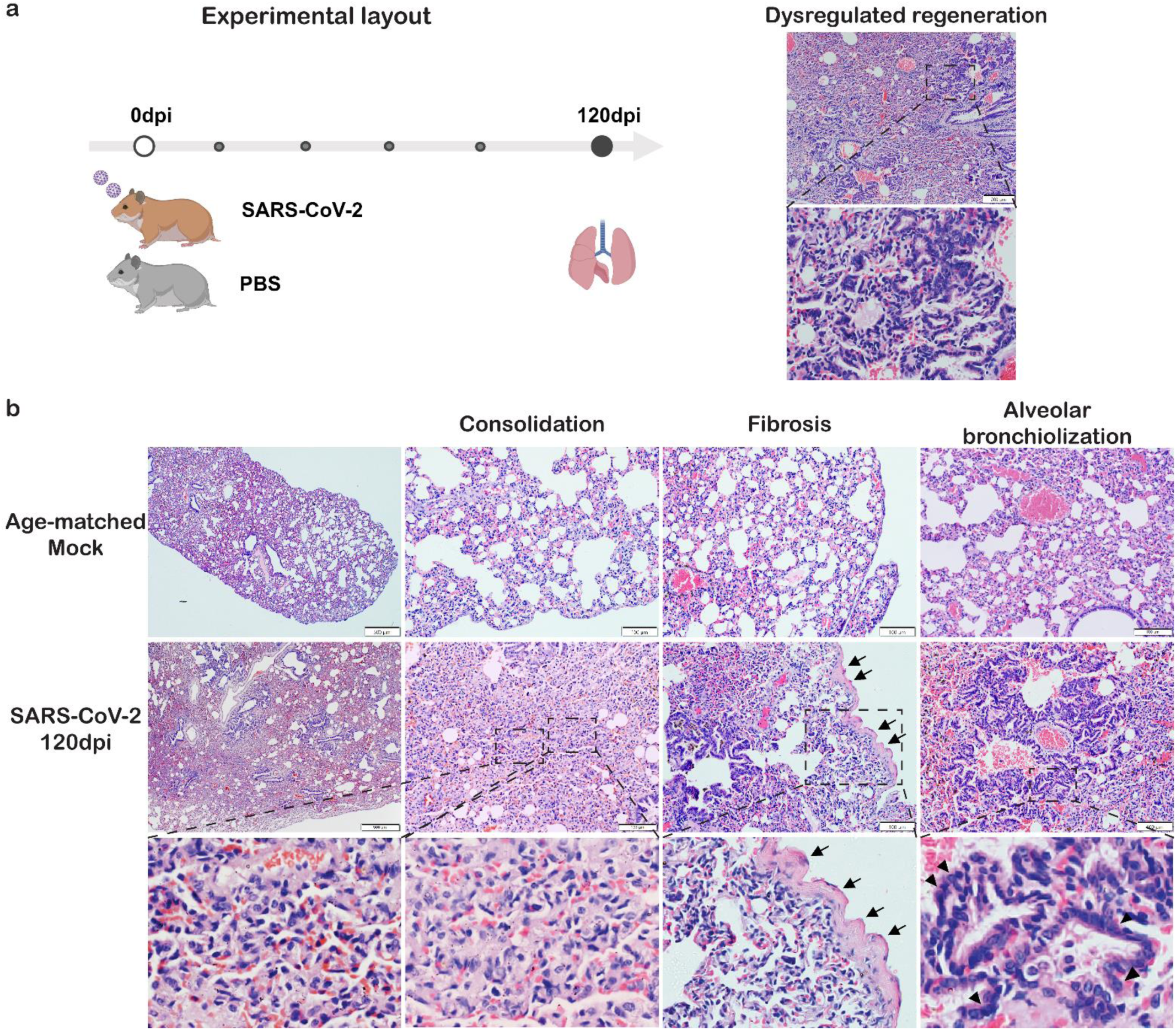
SARS-CoV-2 infection causes persistent abnormal foci of alveolar bronchiolization and fibrosis in hamster lungs. **a.** Experimental layout: 6-8 weeks old hamsters were intranasally inoculated with 10^3^ PFU SARS-CoV-2 wild-type strain HK-13 or equal volume of PBS as mock controls. Lung tissues were collected at 120dpi. Dysregulated regeneration was observed in SARS-CoV-2 infected hamster lungs. Upper H&E image showing lung condensation with blood vessel congestion and multiple abnormal foci. Lower zoomed image showing abnormal foci of alveolar bronchiolizaiton. **b.** Representative H&E images showing pulmonary consolidation, fibrosis and alveolar bronchiolization in SARS-CoV-2 infected hamster lungs at 120dpi. Upper panel: Mock control lung showed normal structure and lung was not inflated. Middle panel: SARS-CoV-2 infected hamsters displayed whole lung condensation and multiple foci from proximal to distal lung. Bottom panel: Higher magnification and zoomed images showing alveolar collapse, alveolar consolidation, pleurae thickening and alveolar bronchiolization. Black open arrows indicated pleurae thickening. Black triangles indicated alveolar epithelial cell hyperplasia. Scale bar=500μm, 200μm, respectively.

We then investigated the progression of lung changes caused by SARS-CoV-2 infection covering different stages of disease including acute (7 dpi), resolving (14 dpi) and chronic phase (42, 84 &120 dpi). Consistent with our previous findings in acute infection (*22*), massive infiltration, severe alveolar consolidation and hyperplastic regeneration were found in SARS-CoV-2 infected hamsters at 7dpi (Fig 2b& Fig S1). At 14dpi, inflammatory infiltration was largely resolved but still detectable; however, abnormal bronchiolization foci forming ribbons and tubules-like structure started to appear (Fig 2b& Fig S1). At 42dpi, diffuse abnormal foci increased and were easily observed in peribronchiolar, perivascular and distal alveolar area (Fig 2b&c). These abnormal foci persisted at 84dpi, and then slightly decreased in size but persisted at 120dpi (Fig 2c&d). Moreover, we found that Omicron BA.5 infected hamsters also displayed persistent abnormal foci and inflammatory infiltration until 120dpi, but at less degree comparing with SARS-CoV-2 wild-type strain HK-13 (Fig S2a), suggesting that chronic lung tissue damages are not SARS-CoV-2 strain-specific.

**Figure 2.**
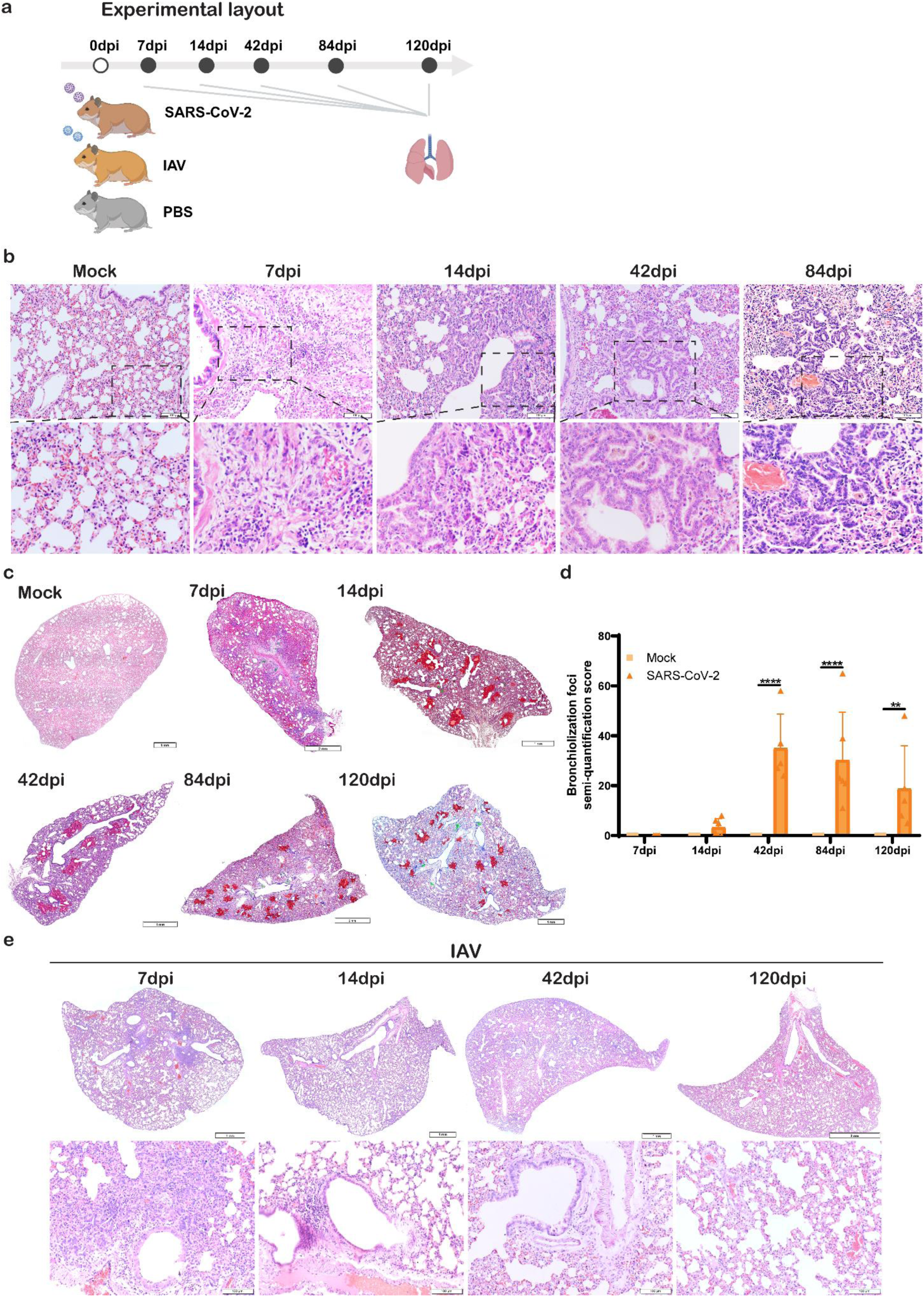
Progression dynamic of SARS-CoV-2 infection caused histopathological changes in hamster lungs. **a.** Experimental layout: 6-8 weeks old hamsters were intranasally inoculated with 10^3^ PFU SARS-CoV-2 wild-type strain HK-13, or 10^5^ PFU mouse-adapted A(H1N1)pdm09 virus (IAV), or equal volume of PBS as mock controls. Lung tissues were collected at 7, 14, 42, 84 and 120dpi. **b.** Representative H&E images showing the prominent histological changes of SARS-CoV-2 infected hamster lungs at 7, 14, 42, 84dpi, and mock control lungs. **c.** Representative full scan H&E image of SARS-CoV-2 infected hamster lung lobe at 7, 14, 42, 84, 120dpi, and mock control. Red highlighted region indicated bronchiolization foci. Green highlighted region indicated infiltration. **d.** Semi-quantification score of bronchiolization foci. n=3-6. **e.** Representative H&E images of IAV infected hamster lungs and age-matched mock control lungs at 7, 14, 42 and 120dpi. Scale bar=2mm, 1mm, 100μm, respectively. Data represented mean±SD. **p<0.01, ****p<0.0001 by Two-way ANOVA.

To verify whether this chronic changes in lung is unique to SARS-CoV-2 infection, hamsters were infected with 10^5^ PFU of mouse-adapted A(H1N1)pdm09 (IAV) for comparison. Consistent with previous findings(*25*), localized perivascular and peribronchiolar infiltration was observed at 7dpi (Fig 2e). Lung inflammatory changes was resolved at 14dpi and returned to normal tissue morphology at 42 and 120dpi without bronchiolization foci or any other histological abnormalities, which indicating that SARS-CoV-2 causes chronic lung tissue damage but not with influenza virus.

### SARS-CoV-2 induced pulmonary fibrosis persisted until 120dpi

Post-COVID pulmonary fibrosis was reported to be the most significant long-term respiratory sequelae (*5*). Lung fibrotic histologyas observed in SARS-CoV-2 infected hamster lungs at 42 and 120dpi, showing as thickened alveolar wall and visceral pleural membrane (Fig 1b). Masson trichrome staining confirmed collagen deposition around bronchi and vessels at 42 and 120dpi (Fig 3a). 71.4% (5/7) at 42dpi and 42.8% (3/7) of hamsters at 120dpi showed pleural fibrosis. Alveolar septa fibrosis is evidenced by increased collagen deposition, which was frequently observed in the lung at 42 and 120dpi. Again, we found that Omicron BA.5 infected hamsters also displayed similar fibrotic changes until 120dpi (Fig S2b). However, no fibrosis was detected at any time points in IAV infected hamster lungs. Real-time RT-qPCR assay demonstrated significant upregulation of the genes involving lung tissue fibrosis in SARS-CoV-2 infected lungs at 120dpi (Fig 3b), including fibroblast growth factor (FGF)1, FGF2 and FGF7. Transforming growth factor-β (TGF-β)1, a well-known major profibrogenic cytokine (*26*), had 10-fold increase after SARS-CoV-2 infection compared with IAV at 42dpi. Imbalanced expression of Matrix metalloproteinases (MMPs) and their specific tissue inhibitors of metalloproteinase (TIMPs) is closely involved in pulmonary fibrosis and fibrolysis (*27*). At 42dpi, MMP2 and MMP9 mRNA level didn’t increase much while TIMP1 and TIMP2 were highly upregulated in SARS-CoV-2 infection than in IAV infection (Fig 3b). At 120dpi, both MMPs and TIMPs were elevated in SARS-CoV-2 infected lungs. Differently, in IAV infection, only slight upregulation of TIMP2 and TIMP3 were observed at 120dpi.

**Figure 3.**
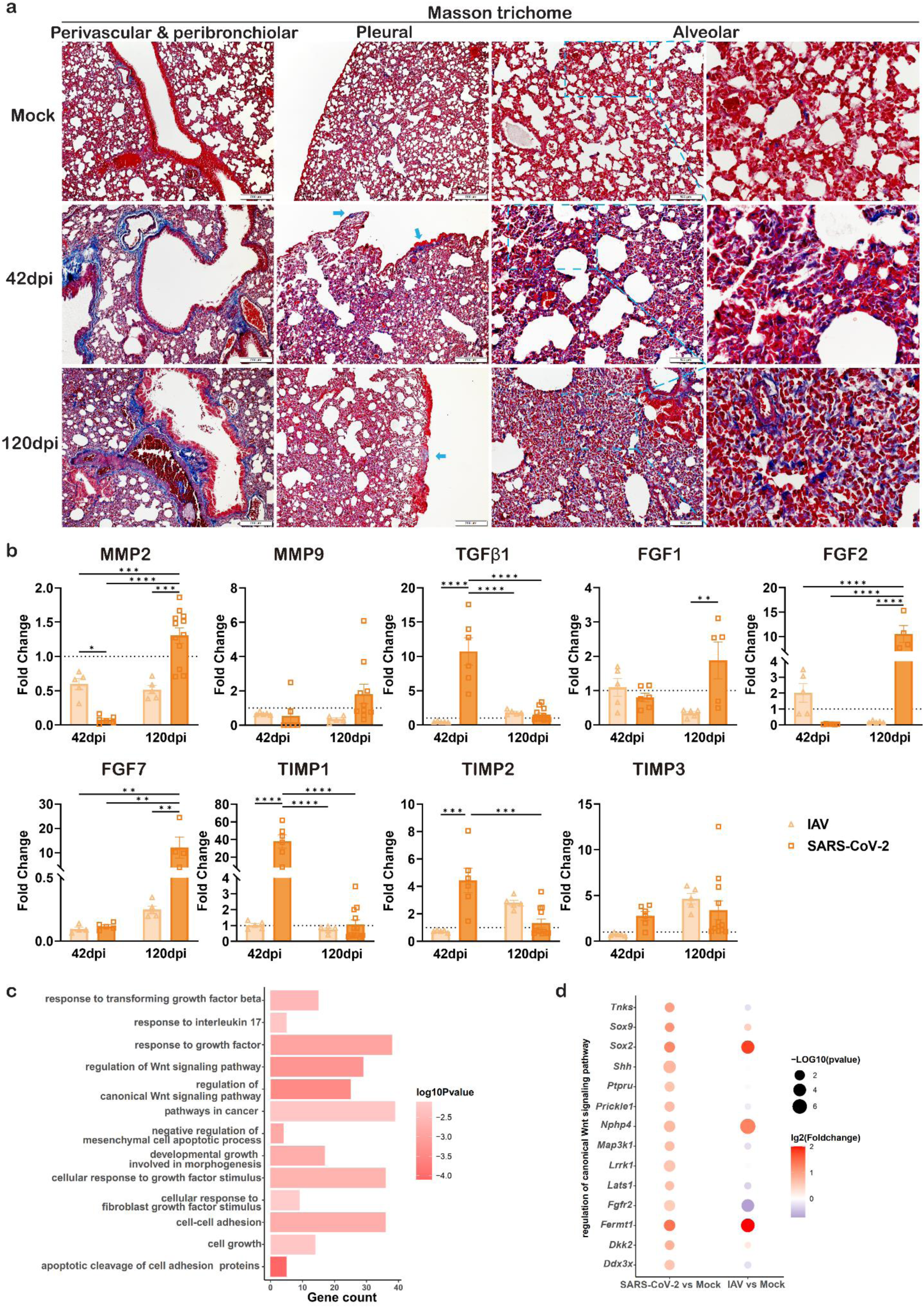
SARS-CoV-2 infection causes persistent fibrosis in hamster lungs. **a.** Representative images of masson trichrome stained lung of mock and SARS-CoV-2 infected hamsters at 42 and 120dpi. Left panel: Representative images showing perivascular and peribronchiolar fibrosis in SARS-CoV-2 infected hamsters. Blue indicated collagen deposition. Second left panel: Representative images showing pleural fibrosis in SARS-CoV-2 infected hamsters. Blue arrows indicated collagen deposition. Right two panels: Representative images and magnified images showing alveolar fibrosis in SARS-CoV-2 infected hamsters. Scale bar=200μm or 100μm. **b.** Relative mRNA expression levels of fibrosis related genes in the lung tissues of SARS-CoV-2 or IAV infected hamsters. n=5-12. Data represents mean±SD. **c.** Upregulated pathways in SARS-CoV-2 infected hamster lungs at 42dpi by RNAseq analysis. **d.** Differential expression of genes in regulation of canonical WNT signaling pathway in SARS-CoV-2 infected hamster lungs at 42dpi by RNAseq analysis. *p<0.05, **p<0.01, ***p<0.001, ****p<0.0001 by One-way ANOVA.

In line with our histological and RT-qPCR findings, RNAseq transcriptomic profiles of hamster lungs 42dpi after SARS-CoV-2 or IAV infection further revealed significant upregulation of genes related to epithelial cell proliferation, fibroblast growth factor signaling and TGFβ signaling pathways in SARS-CoV-2 infected lung tissue compared to mock control (Fig 3c). In addition, genes related to canonical WNT signaling, which stimulates fibroblasts and induces fibrosis, were also upregulated in SARS-CoV-2 infected hamster lung (Fig 3d). This suggests that molecular pathways participating in tissue fibrosis are highly activated at chronic stage in SARS-CoV-2 infected lungs.

### Residual SARS-CoV-2 virus persisted in hamster lungs accompanied with chronic inflammatory responses

In SARS-CoV-2 infected hamsters, cytokine/chemokines mRNA expression of lung tissue increased at 7dpi, then displayed a prolonged upregulation of proinflammatory cytokines and chemokines IL6, TNFα, IFNγ, CXCL10 and CCL3 at 42dpi, only decreased but remained higher than mock at 84dpi (Fig 4a). Notably, chronic inflammatory cytokine IL33 and IL13 were significantly upregulated at 42dpi or 120dpi (Fig 4b). Fibrinogenβ had about 20-fold increase at 120dpi comparing to mock control group. On the contrary, IAV-infected hamsters showed elevated IFNγ, TNFα, CXCL10 and CCL3 mRNA levels at 7dpi and gradually decreased to normal level at 42dpi. These findings indicate that sustained inflammatory responses persisted long after acute phase of SARS-CoV-2 infection.

**Figure 4.**
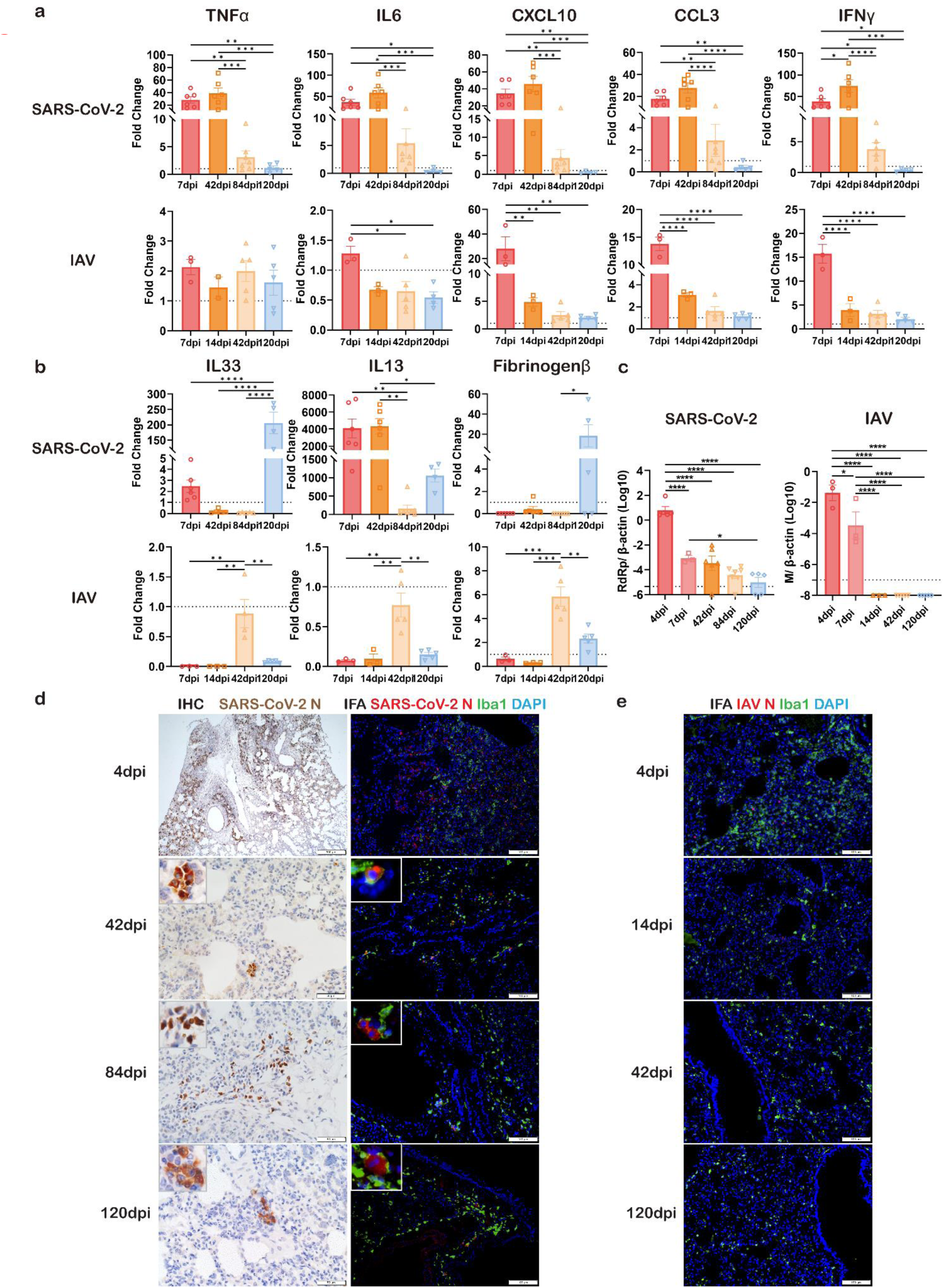
Sustained inflammatory responses and virus persistence in hamster lungs after SARS-CoV-2 infection. **a.** Relative mRNA expression levels of proinflammatory cytokines/chemokines in the lung tissues of SARS-CoV-2 and IAV infected hamsters. n=3-7. **b.** Relative mRNA expression levels of chronic inflammatory mediators in the lung tissues of SARS-CoV-2 and IAV infected hamsters. n=3-7. **c.** Viral load of RdRp (left) and M (right) in the lung tissues at different time points after SARS-CoV-2 or IAV infection. n=3-7. Dashed lines indicated as cut-off line determined by mock controls. **d.** Immunohistochemistry stained SARS2-CoV-2 N protein (left panel) and double immunofluorescence stained N protein (red) and macrophage marker Iba1 (green) (right panel) in SARS-CoV-2 infected hamster lung. Triangles indicated positive cells and magnified images were inserted. **e.** Double immunofluorescence stained IAV N protein (red) and macrophage marker Iba1 (green) in IAV infected hamster lung. Scale bar=100μm. Data represents mean±SD. *p<0.05, **p<0.01, ***p<0.001, ****p<0.0001 by One-way ANOVA.

Now the obvious question is whether observed lung chronic inflammation and dysregulated alveolar regeneration are associated with presence of virus residue in the lung. To this end, we detected SARS-CoV-2 with multiple methods. RdRp gene detection by RT-qPCR showed that viral load peaked at 4dpi, then declined at 7dpi, remained detectable in 5/6 (83.3%) of the hamster lungs at 42dpi, 6/7 (85.7%) at 84dpi and 3/5 (60%) at 120dpi, respectively. As for IAV infected hamster lungs, the results from M gene RT-qPCR showed complete clearance of virus since 14dpi (Fig 4c).

To confirm viral persistence, we performed immunohistochemistry (IHC) and immunofluorescence (IF) staining for SARS-CoV-2 N antigen and IAV N antigen, respectively. As expected, SARS-CoV-2 and IAV infected lungs displayed extensive positive viral antigen expression at 4dpi. Both IHC and IF staining showed SARS-CoV-2 N positive cells in the lung at 42, 84 and 120dpi, while no IAV N antigen could be detected after 14dpi (Fig 4d&e). To identify the cells harboring virus, double IF staining for macrophage marker Ionized calcium binding adapter molecule 1(Iba1) and SARS-CoV-2 N antigen showed double positive cells around terminal bronchiole at 42, 84 and 120dpi. Semi-quantitation indicated that more than 85% of SARS-CoV-2 N+ cells were Iba1 positive, suggesting macrophages are the major reservoir for SARS-CoV-2 in hamster lungs. SARS-CoV-2 N+ cells surrounding bronchiolization foci were also detected, indicating a possible stimulatory role of virus residue to bronchiolization (Fig S3). Taken together, these findings reveal that viral components persist in hamster lungs long after SARS-CoV-2 acute infection, which could be contributing to the chronic inflammatory response and dysregulated alveolar regeneration.

### AT2 cell replenishment in lung regeneration after SARS-CoV-2 infection

Above histopathological and gene transcriptional data indicated dysregulation of tissue repair and lung regeneration after SARS-CoV-2 infection of hamsters. To understand the regeneration process in the lung, we first evaluated the serial changes of AT2 cells. AT1 and AT2 cells are both the primary targets of SARS-CoV-2 virus, while only AT2 cells are considered as progenitors to replenish alveolar epithelium after injury (*10*). Dual IF staining of AT2 cells with SPC and Ki67 at different time points showed dynamic changes of AT2 cells, which decreased significantly at 4dpi, increased at 7dpi and then resumed to the level as of mock control at 14dpi (Fig 5a&b). The percentage of Ki67+ proliferative AT2 cell increased significantly at 4dpi, peaked at 7dp, remained high at 14dpi (Fig 5a&c) which indicated that lung infection activated the undamaged AT2 proliferation function. Further, we found that AT2 proliferation rate returned to base-line level from 42dpi to 120dpi (Fig 5c), indicating no persistent activation of AT2 cells. Notably, AT2 cells were rarely detected in bronchiolization foci from 14 to 120dpi, suggesting that activated AT2 cells are not the origin of the alveolar-bronchiolization.

**Figure 5.**
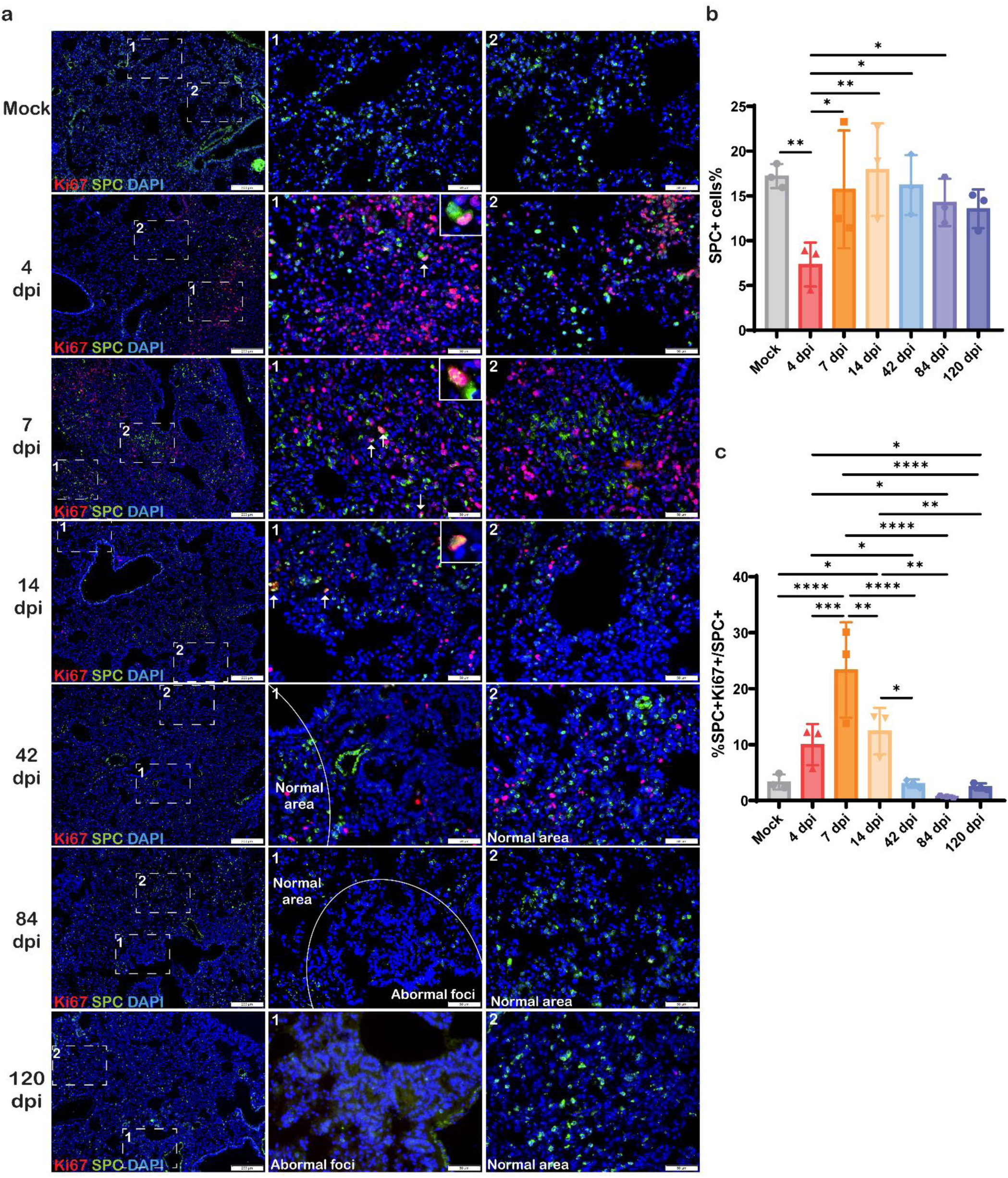
AT2 cells in alveolar regeneration in SARS-CoV-2 infected hamsters. **a.** Representative images of immunofluorescence stained Ki67 (red) and SPC (green) in SARS-CoV-2 infected hamster lungs. White arrows indicated Ki67 and SPC double positive cells and magnified images were inserted. Scale bar=200μm or 50μm. **b.** SPC+ AT2 cells percentage in total cells of mock and SARS-CoV-2 infected hamster lung. n=3. **c.** Proliferative Ki67+ SPC+ AT2 cells percentage in SPC+ AT2 cells of mock and SARS-CoV-2 infected hamster lung. n=3. Data represents mean±SD. *p<0.05, **p<0.01, ***p<0.001, ****p<0.0001 by One-way ANOVA.

### CK14+ basal cells participated airway regeneration and alveolar-bronchiolization after SARS-CoV-2 infection

Next, we systematically investigated CK14+ basal cell’s role in lung regeneration after SARS-CoV-2 infection. We found that CK14+ basal cells increased remarkably in both bronchioles and alveoli at 4 and 7dpi in SARS-CoV-2 infected hamsters, which had a high frequency of Ki67+CK14+ double positivity, while only a few CK14+ basal cells could be seen in the airway of mock control hamster, none in alveoli (Fig 6a&e). This indicates CK14+ basal cells proliferation and migration into alveoli. In alveoli, some CK14+SPC+cells were detected at 7dpi, suggesting CK14+basal cells may contribute to AT2 cells repair to some extent (Fig 6b). At 14 dpi, CK14+ basal cells decreased in bronchiole accompanied with histological evidence of repaired airway epithelium, while they remained frequent in alveoli, clustering in alveoli around bronchioles (Fig 6a). Some of them were co-expressing Ki67 until 42dpi, indicating persistent activation of CK14+ basal cells. Within these cell clusters, CK14+SCGB1A1+ or CK14+Tubulin+ cells were frequently observed at 14dpi (Fig 6c&d), suggesting these CK14+ basal cells differentiated into club cells or ciliated cells in alveoli.

**Figure 6.**
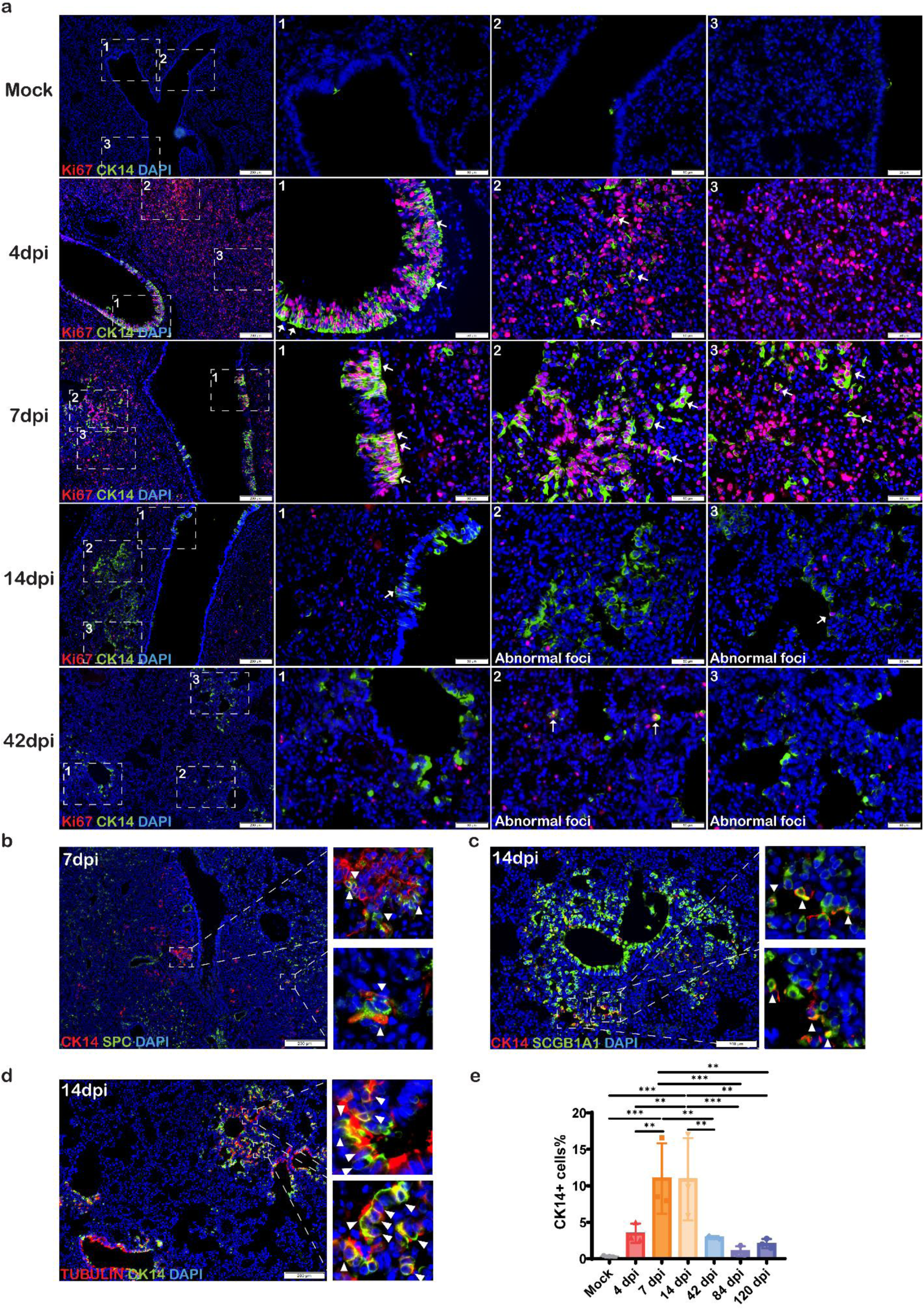
CK14+ basal cells actively proliferated, differentiated in lung regeneration and bronchiolization foci in SARS-CoV-2 infected hamsters. **a.** Representative images of immunofluorescence stained Ki67 (red) and CK14 (green) in SARS-CoV-2 infected hamster lungs. Scale bar=200μm or 50μm. Area 1 indicated bronchioles. Area 2 and 3 indicated alveolar (for infected hamsters). White arrows indicated Ki67 and CK14 double positive cells. **b.** Representative image of immunofluorescence stained CK14 (red) and SPC (green) in SARS-CoV-2 infected hamster lung at 7dpi. White triangles indicated CK14 and SPC double positive cells. Scale bar=200μm. **c.** Representative image of immunofluorescence stained CK14 (red) and SCGB1A1 (green) in SARS-CoV-2 infected hamster lung at 14dpi. White triangle indicated CK14 and SCGB1A1 double positive cells. Scale bar=100μm. **d.** Representative image of immunofluorescence stained CK14 (red) and Tubulin (green) in SARS-CoV-2 infected hamster lung at 14dpi. White triangle indicated CK14 and Tubulin double positive cells. Scale bar=100μm. **e.** CK14+ basal cells percentage in total cells of mock and SARS-CoV-2 infected hamster lung. n=3. Data represents mean±SD. **p<0.01, ***p<0.001 by One-way ANOVA.

To further characterize the cell types in the bronchiolization foci, IF staining was performed for club cells with SCGB1A1 and ciliated cells with Tubulin. SCGB1A1+ club cells and Tubulin+ ciliated cells were only found in the airway epithelium of mock control. In acute SARS-CoV-2 infected lungs, the airway luminal sloughed epithelial cells expressing SCGB1A1 and Tubulin at 4dpi indicating infection caused cell death of club cells and ciliated cells (Fig 7a). Only from 14dpi onward, increased number of SCGB1A1+ cells and Tubulin+ cells were easily observed, forming gradually enlarged foci with ribbon-like or honeycomb-like structure (Fig 7b). Together with the features shown in H&E stained lung sections, we confirmed that SCGB1A1+ club cells and Tubulin+ ciliated cells are the major cellular components of bronchiolization foci in SARS-CoV-2 infected hamster lung. Collectively, these temporal cell type identification and tracing demonstrated that CK14+ basal cells served as major progenitors in the repair of bronchiolar epithelium damaged by SARS-CoV-2 infection, meanwhile they migrated to alveoli and differentiated and formed bronchiolization foci.

**Figure 7.**
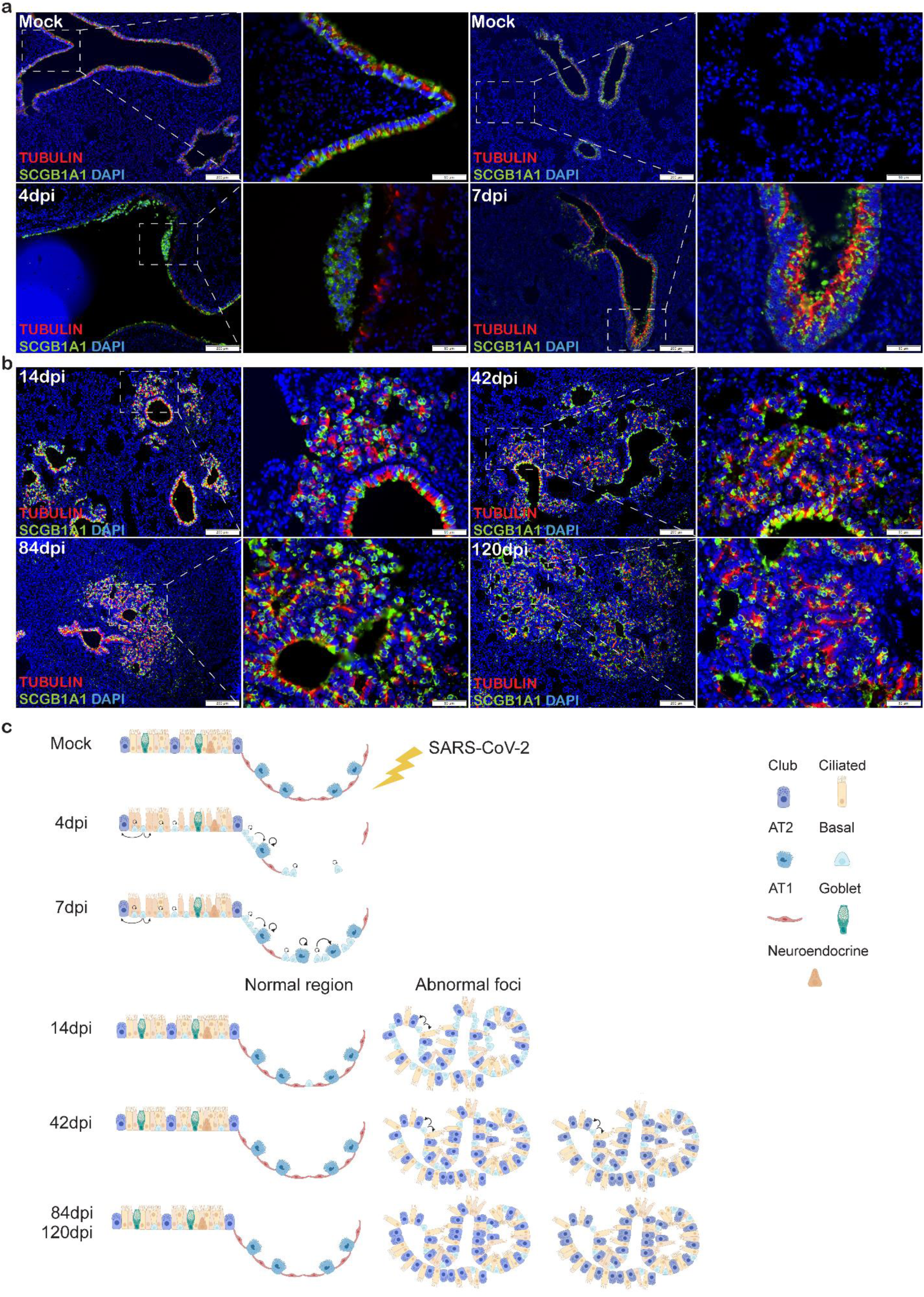
Ciliated cells and club cells consist most of the bronchiolization foci. **a.** Representative image of double immunofluorescence stained Tubulin (red) and SCGB1A1 (green) in mock and SARS-CoV-2 infected hamster lungs at 4 and 7dpi. Scale bar=100μm or 50μm. **b.** Representative image of double immunofluorescence stained Tubulin (red) and SCGB1A1 (green) in SARS-CoV-2 infected hamster lungs at 14, 42, 84 and 120dpi. Scale bar=100μm or 50μm. **c.** Graphic summary of lung regeneration: During acute phase of SARS-CoV-2 infection (4&7dpi), CK14+ basal cells actively contributed to regeneration of brochiolar. In alveolar, AT2 cells were damaged massively due to virus attack. AT2 cells self-renewed, CK14+basal cells migrated and differentiated to AT2 cells for alveolar repair. At 14dpi, most bronchi and part of alveolar resumed to normal structure. CK14+basal cells clustered, differentiated to club cells and ciliated cells, forming abnormal foci of bronchiolization. At 42dpi, CK14+basal cells were still activated. Bronchiolization foci increased in size and number. At 84 and 120dpi, bronchiolization foci persisted and were mostly comprised of ciliated cells and club cells.

### Notch pathway highly activated in chronic phase of SARS-CoV-2 infection

By real time RT-qPCR assay, Notch1 and Notch3 expression levels were found significantly elevated about around 30-40 fold and 80 fold comparing to mock controls at 42dpi and 120pdi, respectively (Fig 8a). Notch2 increased significantly at 42dpi. In addition, Jagged Canonical Notch Ligand (Jag)1, displayed a striking 150-250 fold increase at 42dpi and 120dpi. Notch target genes, Hairy/Enhancer of Split related to YRPW motif (Hey)1 was also highly elevated at 42dpi and 120dpi, while Hairy/Enhancer of Split (Hes)1 decreased. Notch pathway regulatory gene SRY-related HMG-box (SOX)2 increased at 120dpi. On the contrary, IAV infected hamsters only showed slight increase of Jag1, no other detectable changes at 42 or 120dpi.

**Figure 8.**
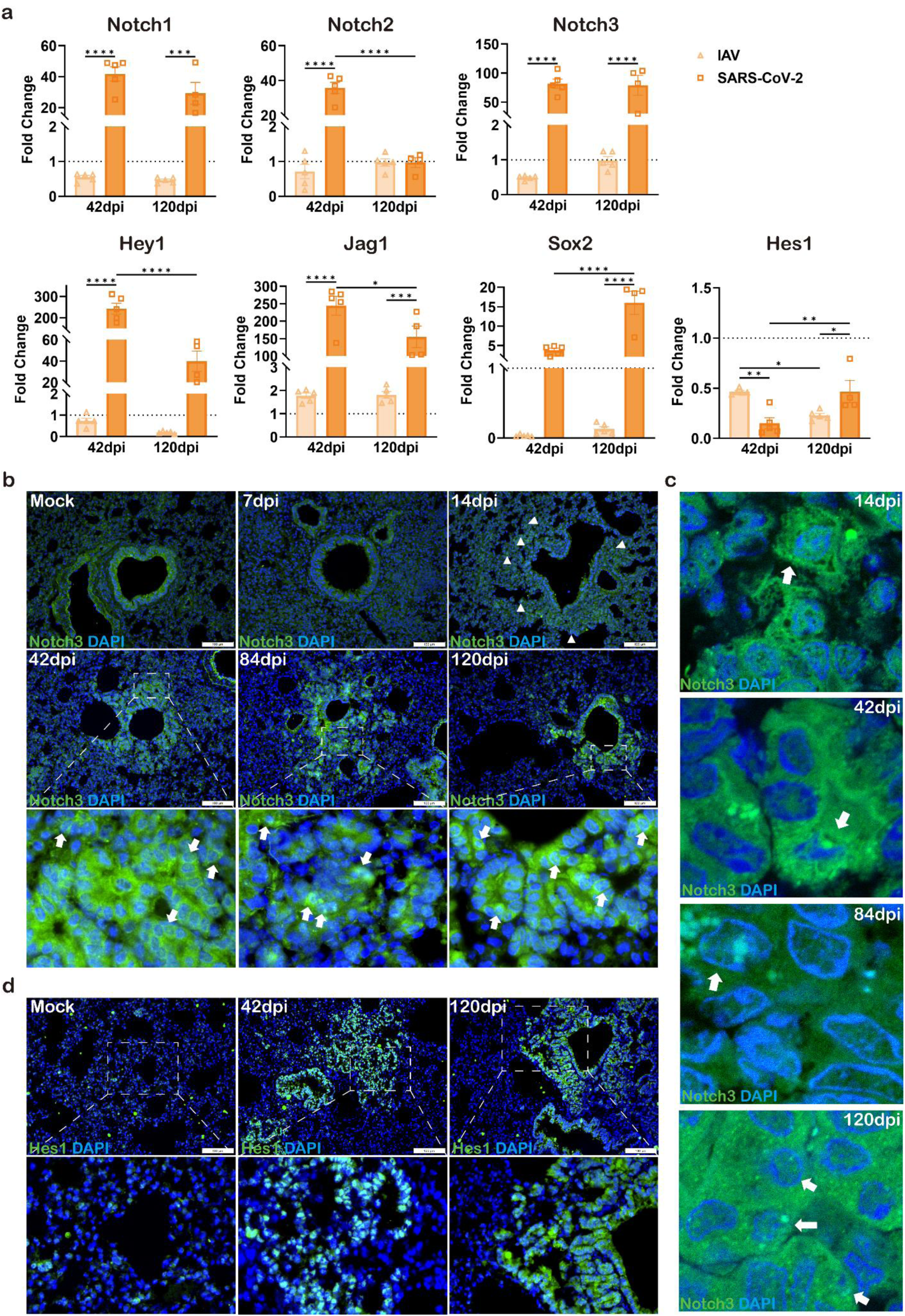
Highly elevated Notch signaling long after SARS-CoV-2 infection in hamster lungs. **a.** Relative mRNA expression levels of Notch signaling related genes in the lung tissues in SARS-CoV-2 and IAV infected hamsters. n=4-5. Data represents mean±SD. **b.** Representative images of immunofluorescence stained Notch3 (green) in mock and SARS-CoV-2 infected hamster lung at 7, 14, 42, 84 and 120dpi. Scale bar=100μm. White triangles indicated Notch3 positive cells. White arrows indicated Notch3 signals inside nuclei. **c.** Representative magnified images of immunofluorescence stained Notch3 (green) captured by confocal microscopy in SARS-CoV-2 infected hamster lung at 14, 42, 84 and 120dpi. White arrows indicated Notch3 signals inside nuclei. **d.** Representative images of immunofluorescence stained Hes1 (green) in mock and SARS-CoV-2 infected hamster lung at 42 and 120dpi. Scale bar=100μm. White arrows indicated Notch3 positive cells. *p<0.05, **p<0.01, ****p<0.0001 by One-way ANOVA.

Moreover, detection of Notch3 protein by IF staining showed sporadic Notch3 positive cells appeared in alveolar area at 14dpi, while intensive staining of Notch3 positive cells clustered in the lung section at 42, 84 and 120dpi (Fig 8b). The distribution of Notch3 high-expression cells matches the abnormal bronchiolization foci microscopically. Notch3 nuclear localization by IF staining in a portion of the cells further indicates activation and nuclear translocation of Notch3 (Fig 8c). Moreover, IF stained Hes1 protein was also detected with high expression in cells clustered in the abnormal bronchiolization foci of SARS-CoV-2 infected hamsters at 42 and 120dpi (Fig 8d). Collectively, our data indicate continuous and persistent activation of Notch pathway in the chronic phase of SARS-CoV-2 lung infection.

### Notch pathway transcriptional changes and upregulation of cancer related genes spatially overlapping alveolar-bronchiolization region in the lung

Alveolar-bronchiolization is known as a potential precursor lesion of lung cancer (*28*). On the other hand, abnormal expression of Notch signaling pathway is frequently observed in lung cancer cases (*29*). To investigate if the potentially Notch dysregulation induced bronchiolization caused by SARS-CoV-2 infection is possible risk factor for lung cancer, we conducted spatial transcriptomic experiment using hamster lung samples collected at 120dpi, comparing the transcriptomics profile of the histologically abnormal lung regions with that of the normal regions. Six tissue regions were identified and selected to represent abnormal (red circled) or normal (black circled) alveolar structure based on histological features observed in the H&E section (Fig 9a). Firstly, cell type signature genes for lung epithelium (*8, 10*), including AT1, AT2, ciliated cells and club cells were applied for spatial mapping. For instance, *Ccdc39*, which is highly expressed in ciliated cells (*30*) was abundantly detected in abnormal circle regions (Fig 9b). *Foxj1*, which induces basal cells differentiation to ciliated cells, was detected only in abnormal circle regions (Fig 9b&S4). We observed higher expression of ciliated cell and club cell marker genes, together with lower expression of AT1 and AT2 cell marker genes in abnormal region comparing with normal region (Fig 9c). These data from transcriptional level proved the aberrant cellular composition in the alveolar abnormal regions. Secondly, pathway enrichment analysis of the DEGs between the normal and abnormal circles revealed the upregulation of genes associated with positive-regulation of cell growth, position-regulation of GTPase activity, Wnt signaling pathway, NF-kappaB and ERBB signaling pathways in the abnormal regions (Fig 9d). Moreover, genes associated with positive regulation of Notch signaling pathway (GO:0045747) were also upregulated in the abnormal regions (Fig 7e). To be noted, significant upregulation of several pro-tumor genes or genes positively regulating cell cycles, including tubulin beta 4B class IVB (*Tubb4b*), syntaxin binding protein 4 (*Stxbp4*), growth factor receptor bound protein 14 (*Grb14*), Myeloid leukemia factor 1 (*Mlf1*), Mucin 1 (*Muc1*) and P53 Apoptosis Effector Related To PMP22 (*Perp*) were detected in abnormal regions (Fig 9f). Upregulation of these genes was observed in lung cancer tissues and some genes were demonstrated to drive lung tumor growth (*31–36*). This indicate the abnormal regions are possible to be precursor of lung cancer. Taken together, spatial-specific transcriptomic data provide further evidence supporting dysregulated lung regeneration long after SARS-CoV-2 infection. More importantly, our data suggests the possibility of increased risk of lung cancer in long COVID.

**Figure 9.**
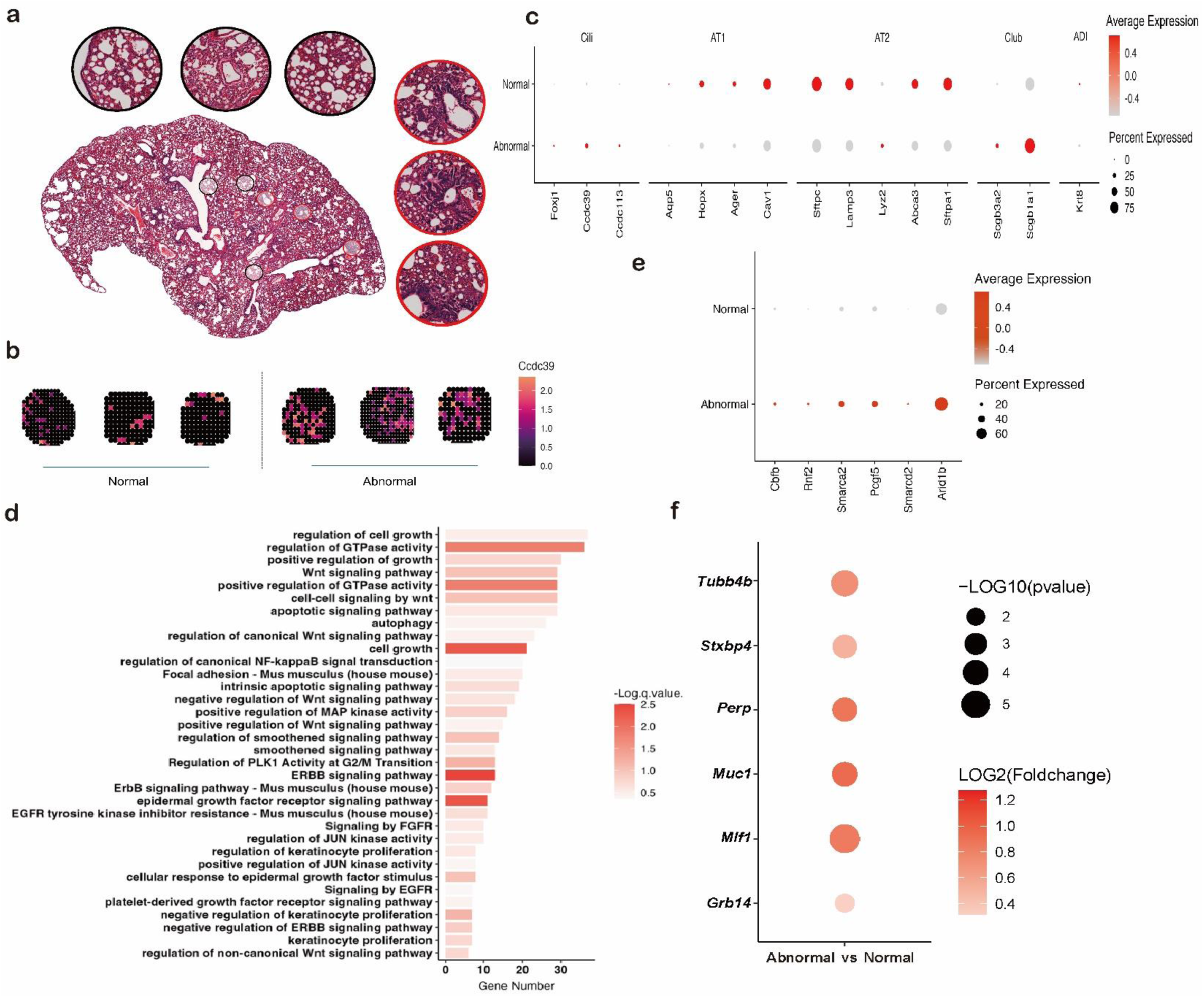
Spatial transcriptomics of SARS-CoV-2 infected hamster lung. **a.** Six tissue regions representing abnormal (red circled) or normal (black circled) alveolar structures based on histological features shown in the H&E section. **b.** *Ccdc39* gene expression in selected normal (left) and abnormal (right) tissue regions. **c.** Differential expression of genes expressed on ciliated cells, AT1 cells, AT2 cells, club cells and ADI cells in selected normal and abnormal tissue regions. **d.** Enriched pathways based on the up-regulated genes in abnormal tissue regions. **e.** Differential expression of genes associated with positive regulation of Notch signaling pathway in selected normal and abnormal tissue regions. **f.** Differential expression of genes related with cancer in selected normal and abnormal tissue regions.

## Discussion

A substantial number of patients with physical and/or mental sequelae long after SARS-CoV-2 acute infection becomes a worldwide health problem drawing the attention of life-science and medical research society. Patients with respiratory PACS, often complain on shortness of breath or dyspnea, which may or may not accompanied with radiologically detectable lung abnormalities including inflammation, ground-glass opacities or fibrotic changes (*4, 5*). Investigations of lung histopathological changes and underlying molecular mechanisms are urgently needed. In this study, we demonstrated chronic inflammatory damages with residual virus residing in macrophages in SARS-CoV-2 infected hamster lung beyond 120 dpi. Excessive proinflammatory cytokines responses caused by SARS-CoV-2 acute infection incompletely resolved and sustained up-to 42 days when chronic inflammatory mediator increased with lung fibrotic lesions. Alveoli regeneration process following SARS-CoV-2 infection disrupted by dysregulated CK14+ basal cells migration and differentiation into SCGB1A1+ club cells and Tubulin+ ciliated cells, which develop into persistent alveolar-bronchiolization. Further, our data indicated that continuous upregulation of Notch pathway and related genes contributes to the abnormal lung tissue regeneration.

SARS-CoV-2 acute infection hamster model simulated perfectly the lung pathology in COVID-19 patients, inflammatory tissue damages peak at 4dpi and start to resolve at 7dpi and recovered at 14dpi (*22, 24*). However, in our current longitudinal study, we found the expression levels of proinflammatory cytokines and chemokines IL6, TNFα, CXCL10 and CCL3 remained as high as 7dpi at 42dpi, back down but remain above baseline level until 120dpi; while IL33, IL13 and Fibrinogenβ were highly upregulated at 120dpi, indicating a state of chronic inflammation. As in COVID-19 patients, ours and many others previous animal study showed that SARS-CoV-2 virus was cleared without viral shedding from oral or fecal route when the host mounted effective adaptive immunity one to two weeks after infection (*24, 37*). To elucidate whether there is residual SARS-CoV-2 in the lung, we explored different detection methods and found viral RdRp gene and nucleoprotein in hamster lung tissues by RT-qPCR, IHC and IF, respectively. This finding is consistent with accumulating clinical reports showing virus residues in various human organs long after acute disease (*38*). We further proved that the virus mainly resides in lung macrophages by IF staining of viral N protein and macrophage marker Iba1 (more than 85%), while a small number of N **^+^**/Iba1^-^ cells were not identified. In human studies, SARS-CoV-2 N protein was detected in macrophages in secondary lymphoid tissue autopsy by immunostaining and SARS-CoV-2 RNA was found in CD68+ macrophages in lung tissues by RNA-FISH and consecutive IHC (*39, 40*). Another longitudinal study in cynomolgus macaques showed that replication-competent virus in bronchoalveolar lavage fluid macrophages 6 months post SARS-CoV-2 infection without evidence of infectious virus produced (*41*). Though we did not have evidence whether the virus is capable of replication in hamster macrophages in the current study, our findings together with other reports clearly demonstrated evidence of SARS-CoV-2 residue in the lung tissue. Viral persistence in macrophages is very likely to be an inflammation inducer and contribute to sustained inflammatory responses in the lung. Persistent viral components in macrophages could induce cytokine responses through engagement with host pattern-recognition receptors. Repeated recognition of persistent stimuli might lead to effector activity, exhaustion and altered differentiation of virus-specific T cells and B cells, which will further result in tissue damage or cytopathology (*38*). SARS-CoV-2 proteins are documented to dysregulate host signaling pathways activity through disrupting host metabolic, genetic and epigenetic regulations (*42*), which may also contribute to chronic inflammation. In line with studies in human and hamsters (*6, 43*), peribronchiolar and perivascular cell infiltration was observed in our SARS-CoV-2 infected hamsters from 14 to 120dpi. Among the infiltrating immune cells, we found many Iba1+ macrophages, suggesting the active roles of macrophages in prolonged pulmonary inflammation in SARS-CoV-2 infected hamsters.

Prolonged lung inflammation and presence of viral components are undoubtedly associated with dysregulation of lung tissue repair and regeneration. Lung tissue transcriptomics analysis indicated Fibrosis growth factor and TGFβ signaling pathway related genes were significantly upregulated in SARS-CoV-2 infected lungs at 42dpi. By RT-qPCR, we also detected significant upregulation of TGF-β1, TIMP1 and TIMP2 at 42dpi. The expression of FGF1, FGF2 and FGF7 were significantly high at 120dpi. These molecular changes indicate the activation of pulmonary fibrotic process. Histologically, we found that widespread lung tissue fibrosis, collagen deposition in the visceral pleural membrane, lung interstitial connective tissue and alveolar septa. This is highly consistent with clinical reports in human (*5, 6*).

The most remarkable histological abnormality found in SARS-CoV-2 infected hamster lung at 42-120dpi was the multifocal ribbon-like or honeycomb-like epithelial structures distributed in peribronchiolar, perivascular and distal lung regions. This is in line with previous studies showing bronchiolar epithelium–like cell hyperplasia, characterized as alveolar-bronchiolization, in hamsters recovered from SARS-CoV-2 infection (*8, 43, 44*). The diffuse alveolar-bronchiolization occurred after SARS-CoV-2 infection but not in A(H1N1)pdm09 influenza virus infection. And there is no evidence of residual virus, nor chronic inflammatory lesion after influenza infection. Our hypothesis is that the destruction of multiple types of lung cells, from airway to alveoli, in SARS-CoV-2 acute infection induces excessive inflammatory responses. The inflammation response persists and creates a molecular niche, which dysregulates progenitor cells proliferation and differentiation affecting normal tissue regeneration process. To prove this hypothesis, our analysis of serial lung tissues showed that upon the severe loss of AT2 cells at 4dpi, AT2 cells were activated and proliferated to replenish the damage alveoli. This is in line with the role of AT2 cells’ self-renewal in alveoli regeneration and homeostasis (*11*). From 7dpi onward, AT2 cell number returned to base-line, suggesting AT2 cells regeneration remained relatively unaffected. We did not evaluate AT2 cells’ function as a progenitor to replenish the lost AT1 cells. But Heydemann and colleagues (*8*) reported that in SARS-CoV-2 infection hamster lung, the process of AT2 regeneration for AT1 cells was impaired with a prolonged state of alveolar differentiation intermediate (ADI) cells expressing CK8+. They suggested that multipotent CK14+ basal cell activation and migration into alveoli to facilitate alveolar regeneration, however, the longest observation time in their study was 14dpi. Our data indicates that CK14+ cells played limited roles in AT2 cell replenishment when they migrated out of the airway in hamster lung, though we did see a few CK14+ basal cells co-expressing SPC at the early time of tissue repair.

It is widely accepted now that multiple types of airway progenitor cells contribute to alveolar regeneration after lung damage. Studies in human and mice showed CK5+ basal cells involving in alveolar repair (*12, 19, 45*). To this end, CK5+ basal cells were activated and proliferated to form proliferating cell clusters termed “keratin 5 pods”, a common feature of epithelial remodeling after alveoli damage (*12, 19, 45*). However, consistent with a previous report on hamster model (*8*), we did not detect much CK5+ cells in mock or infected hamster lungs either in bronchiolar or alveolar structures. We assumed that CK14+ basal cells in hamsters play critical role in lung regeneration, which could be equivalent to CK5+ basal cells in human and mice. At 4dpi, CK14+ basal cells were activated and proliferated to replenish the lost bronchiolar epithelium, which was restored completely at 14dpi as shown by H&E tissue sections. Like CK5+ basal cells, CK14+ basal cell progenitors are multipotent. We found CK14+ basal cells migrated into the alveoli, where most of them continued to be proliferative and differentiated to club cells and ciliated cell instead of differentiating into AT2 or AT1 cells. So, there is a possibility that the function of CK14+ basal cells proliferation and differentiation was deregulated under these circumstances, resulted in the formation of alveolar-bronchiolization, though the detailed developmental trajectory needs further clarification.

Basal cells self-renewal and differentiation as progenitor to other cells are regulated by complex molecular signaling pathways (*14*). Notch signaling pathways play important roles in regulating of basal cell differentiation (*14, 46, 47*). Notch3 activation primes basal cells to suprabasal cells (*47*), which subsequently differentiate to club cells or goblet cells through Notch1/2 and Hey signaling activation (*47*). It has been previously showed that basal cell can directly differentiate to ciliated cells during lung repair through Notch2 activation and Sox2 upregulation (*48*). Sustained Notch activation promotes basal cells differentiation toward club cell lineage and inhibits basal cell trans-differentiation to alveolar epithelia cells (*12, 14*). In SARS-CoV-2 infected hamster lungs, elevated expression of Notch1, Notch3, Hey1, Jag1 and Sox2 at 120dpi is in line with these previous reports. Further, spatial transcriptomic data showed that genes associated with positive regulation of Notch signaling pathways were upregulated in the abnormal foci at 120dpi. Consistent with our findings, Notch signaling was upregulated at transcriptional level in SARS-CoV-2 infected juvenile macaque lungs at 14dpi, however no lung pathological was described (*49*), which may be due to the short observation period. Continuous Notch activation was also observed in influenza virus infection in mice (*12*), which implicated the association of Notch pathway activation with abnormal lung repair and fibrosis. Notch activation is required for lung tissue regeneration after infection or injury, while imbalanced Notch signaling may lead to dysregulated lung regeneration. Identification and characterization of specific changes in Notch pathways involving dysregulated lung regeneration could be of potential value for developing therapeutic approaches against chronic pulmonary abnormalities after severe injury or virus infection.

In relation to pulmonary sequela of SARS-CoV-2 infection, besides causing the respiratory symptoms, our data suggest probable risk of lung carcinoma. Firstly, pulmonary fibrosis is well accepted as a risk factor for developing lung cancer since they share many cellular and molecular pathways (*50*). Secondly, alveolar-bronchiolization, potential precursor lesion of lung cancer, was detected in up to 12% of non-small cell lung cancer resection specimens (*28*). Thirdly, upregulation of some genes, which were previously reported in lung tumor tissues and drive tumor growth, was detected. Notch pathway is vital in cell fate determinations and is actively involved in tumor cell initiation, proliferation, differentiation and apoptosis (*29, 51*). Notch1 is reported to promote tumor initiation as an oncogene (*29, 52*). Though we failed to detect Notch1 protein level due to limited reagents against hamster, we demonstrated significant increase of Notch1 genes at 42 and 120dpi. Amplification of Sox2 drives bronchial dysplasia and is an early and consistent event in lung squamous cell carcinoma (SCC) (*53*). We found Sox2 gene expression was significantly elevated in SARS-CoV-2 infected hamster lung at 120dpi. Moreover, our spatial transcriptomic data showed significant upregulation of several pro-tumor genes in the abnormal bronchiolization region. *Stxbp4* is demonstrated to drive lung SCC growth and predictive marker associated with worse prognosis in lung SCC (*35*). *Tubb4b* overexpression is reported in several cancers (*36*). Recent studies implicated that *Tubb4b* is essential for cancer stem cell niche maintenance via cooperation with Ephrin-B1 and *Tubb4b* promotes tumor growth and metastasis through lung cancer cells and monocyte crosstalk (*36, 54*). Studies also demonstrate upregulation of *Mlf1* in lung SCC (*31*), elevation of *Perp* (*32*) and *Grb14* in lung adenocarcinoma (*34*), and increased *Muc1* expression in lung cancer and precancerous lesions (*33*). Taken together, our data suggest increased risk of lung carcinoma after SARS-CoV-2 infection, especially in those with lung abnormalities. Whether SARS-CoV-2 caused chronic lung tissue histological changes are predisposed for malignancy worth tight monitoring, long-term surveillance and further investigation are warranted.

In summary, our study provides evidence of SARS-CoV-2 acute infection caused persistent chronic inflammatory and fibrotic lesion with residual viral components lingering in the host, together with the multiple foci of alveolar-bronchiolization in the lung. This could be the major cause for sustained respiratory symptoms. Aberrant CK14+ basal cells activation and differentiation, which is potentially mediated by continuous hyperactive Notch signaling, contributed to dysregulated lung regeneration. Moreover, there’s potential increased risk of lung carcinoma in Long-COVID patients with chronic lung changes, which requires further investigation and possible surveillance. Our findings provided important information for further exploration of the potential therapeutic approaches and the management of long-COVID-19.

## Methods

### Virus, Cell lines and Biosafety

SARS-CoV-2 wild-type strain HK-13 (GenBank accession no. MT835140), and BA.5 strain (GISAID accession number: EPI_ISL_13777658) were isolated from laboratory-confirmed COVID-19 patients, propagated and titrated in African green monkey kidney cells overexpressing TMPRSS2 protein (Vero E6-TMPRSS2) (*25*). Mouse adapted strain of A(H1N1)pdm09 (A/HK/415742/2009) was propagated in eggs and titrated in Madin-Darby canine kidney cells (MDCK)(*25*). Experiments involving infectious SARS-CoV-2 were performed following the approved standard operating procedures of the Biosafety Level 3 facility.

### Animals

Syrian golden hamsters, 6–8 weeks old, were obtained from the Centre for Comparative Medicine Research, HKU. All the animal experimental procedures were approved by the Committee on the Use of Live Animals in Teaching and Research of HKU (CULATR #5783-21 and 6093-22) and the Institutional Review Board of Beijing Genomics Institute, Shenzhen, China (BGI-IRB A21031-T2).

### Virus challenge in hamsters

Hamsters were inoculated intranasally with 10^3^ PFU SARS-CoV-2 or 10^5^ PFU mouse-adapted A(H1N1)pdm09 virus in 50μl volume per animal under anesthesia of ketamine (150mg/kg) and xylazine (10mg/kg) as we previously reported (*25*). Mock-infection controls were given the same volume of PBS. At 4, 7, 14, 42, 84 or 120dpi, hamsters were euthanized by intraperitoneal injection of pentobarbital sodium (200mg/kg). Blood and lung tissues were harvested for analysis.

### RNA extraction and RT–qPCR

Total RNA extracted from homogenized lung tissues was reverse transcribed into cDNA using MiniBEST Universal RNA Extraction Kit (Takara) and PrimeScriptTM RT reagent kit (Takara). RT-qPCR was performed using QuantiNova Probe PCR Kit (Qiagen) or SYBR Premix Ex Taq II Kit (Takara) with gene-specific primers (Supplementary Table 1). The expression of cytokine/chemokine and Notch signaling pathway/Fibrosis related genes was analyzed by ΔΔCt method, using age-matched naïve samples as baseline.

### Histopathology, immunohistochemistry and immunofluorescence staining

Formalin-fixed, paraffin-embedded lung tissues were processed into 4µm sections and stained with hematoxylin and eosin (H&E) for histopathological examination. Viral antigens and cellular proteins were detected by immunohistochemistry or immunofluorescence with specific antibodies: home-made rabbit/mouse anti-SARS-CoV-2 nucleocapsid protein antibody (*25*), home-made mouse anti-H1N1 nucleocapsid protein antibody (*25*), anti-Ki67 antibody (556003, BD; ab15580, abcam), anti-β tubulin-IV antibody (T7941, Sigma), anti-CK14 antibody (PA516722, Invitrogen; MA5-11599, Invitrogen), anti-SCGB1A1 antibody (10490, Proteintec), anti-SPC antibody (ab3786, Millipore), anti-Iba1 antibody (ab178846, abcam), anti-Notch3 antibody (MBS242006, mybiosource) and anti-Hes1 antibody (LS-B2211 LSBio). Fibrosis staining was performed using NovaUltra Massion Trichrome Stain Kit (IW3006, IHC World) according to the manufacturer’s instructions.

### Digital image analysis

All tissue sections were examined under light or fluorescence microscopy in a blinded fashion. Whole slide images were captured with Akoya Vectra Polaris Scanner at 200× magnification (Akoya Bioscience, USA). Sectional Images were captured by Olympus BX53 Light Microscope (Olympus Life Science, USA). Confocal images were captured with Zeiss LSM900 with AiryScan 2 with 40× and 63× oil objectives. Bronchiolization foci were annotated with QuPath and semi-quantification scoring was manually performed (Figure S5). For quantification of specific cell marker positive cells, 8-12 images of immunofluorescence-stained lung sections were randomly captured in the whole lung. Exported TIFF format files were analyzed with QuPath (Version 0.4.4).

### Bulk RNA-seq experiment and data analysis

Total RNA from lung tissue cells for hamsters was isolated using NucleoSpin RNA Kit (740955.250, MACHEREY-NAGEL, Duren, Germany). cDNA libraries were prepared by KAPA mRNA HyperPrep Kit following the standard protocol. The libraries were denatured and diluted to optimal concentration. Illumina NovaSeq 6000 was used for Pair-End 151bp sequencing. Raw sequencing data were filtered with fastp v0.20.1 to remove adapters and low-quality reads. The alignment file was used for assembling transcripts, estimating their abundances, and detecting differential expression of genes. The genes expression level was quantified by HISAT2 v2.2.0 (*55*) (Reference genome: BCM_Maur_2.0 & grcm38/release_M25). Principal components analysis (PCA) was conducted in R v4.0. Differentially expressed genes were determined based on counts using DESeq2 v3.15 (*56*) based on the following thresholds: |log2FC| > 1 and FDR value <0.05. Metascape (*57*) was used to conduct enrichment analysis involving Gene Ontology (GO) and the Kyoto Encyclopedia of Genes and Genomes Pathway (KEGG) for the DEGs. The reference species was *Mesocricetus auratus*.

### Spatial transcriptomics experiment and analysis

Formalin-fixed, paraffin-embedding (FFPE) lung tissues were collected for Stereo-seq experiment. Library preparation was performed using the Stereo-seq Transcriptomics Set for FFPE (BGI-Shenzhen, China) according to the manufacturer’s protocol. The Stereo-seq chip was 1cm×1cm in size, which contained capture spots with a diameter of 220nm and a spot-to-spot distance of 500nm (*58*). The capture probes contained random primers to capture RNAs. In brief, 5μm FFPE lung tissue section was cut, fully spread, and placed on the Stereo-seq chip. One adjacent tissue section was cut for H&E staining. After dewaxing and decrosslinking reaction, the tissue section was permeabilized to release RNA for in situ hybridization. After RNA capture, reverse transcription was performed and cDNA was later released. The cDNA was then purified and amplified with PCR mix. The PCR products were used to prepare library, which was sequenced by an MGI DNBSEQ T series sequencer (MGI-TECH, China). Raw sequencing data were processed using the SAW pipeline (https://github.com/BGIResearch/SAW). High-quality reads were annotated to generate the spatial gene expression matrix by handleBam (*59*). Seurat (v4.9.9) was used for further quality control, SCT normalization, dimensionality reduction, clustering, and identification of gene expression patterns (*60*). The spatial transcriptomic data was analyzed with a unit of bin100 (49.72μm×49.72μm).

### Data sharing

The spatial transcriptomic data and Bulk RNAseq transcriptomic data supporting this study have been deposited into CNSA (CNGB Sequence Archive) of CNGBdb (https://db.cngb.org/cnsa/) under the project number STT0000082 and CNP0005559.

### Statistical analysis

All data were analyzed with Prism 8.0 (GraphPad Software Inc). Statistical comparison among different groups were performed using One-way or Two-way ANOVA. p<0.05 was considered statistically significant in all figures.

## Author contribution

CL, NX, WS, AH-CL, FL, JF-WC, K-YY, HC and AJ-XZ had roles in the study design, data collection, data analysis, data interpretation, and writing of the manuscript. ZY, YC, JC and AC-YL had roles in the experiments, data collection, data analysis. ZO, WS, XC and PR conducted the Stereo-seq experiment and analysis. HC had roles in data interpretation and writing of the manuscript. All authors reviewed and approved the final version of the manuscript.

## Acknowledgement

We thank China National GeneBank for providing sequencing services and STOmics Cloud (https://cloud.stomics.tech) for their support in Stereo-seq data analysis. We acknowledge the assistance from the University of Hong Kong Li Ka Shing Faculty of Medicine Faculty Core Facility. This study was partly supported by funding from the Collaborative Research Fund (C7060-21G) and Theme-Based Research Scheme (T11-709/21-N), the Research Grants Council of the Hong Kong Special Administrative Region; the National Natural Science Foundation of China General Program (82272337); the Health and Medical Research Fund (20190572), the Food and Health Bureau, The Government of the Hong Kong Special Administrative Region; Health@InnoHK, Innovation and Technology Commission, the Government of the Hong Kong Special Administrative Region; Partnership Programme of Enhancing Laboratory Surveillance and Investigation of Emerging Infectious Diseases and Antimicrobial Resistance for the Department of Health of the Hong Kong Special Administrative Region Government; Sanming Project of Medicine in Shenzhen, China (SZSM201911014); the High Level-Hospital Program, Health Commission of Guangdong Province, China; National Key Research and Development Program of China(projects 2021YFC0866100 and 2023YFC3041600); Emergency Collaborative Project of Guangzhou Laboratory (EKPG22-01); the Major Science and Technology Program of Hainan Province (ZDKJ202003); the research project of Hainan Academician Innovation Platform (YSPTZX202004); the University of Hong Kong Outstanding Young Researcher Award; and the University of Hong Kong Research Output Prize (Li Ka Shing Faculty of Medicine); and donations from May Tam Mak Mei Yin, Richard Yu and Carol Yu, the Shaw Foundation Hong Kong, Michael Seak-Kan Tong, Lee Wan Keung Charity Foundation Limited, Providence Foundation Limited (in memory of the late Lui Hac-Minh), Hong Kong Sanatorium and Hospital, Hui Ming, Hui Hoy and Chow Sin Lan Charity Fund Limited, The Chen Wai Wai Vivien Foundation Limited, Chan Yin Chuen Memorial Charitable Foundation, Marina Man-Wai Lee, the Hong Kong Hainan Commercial Association South China Microbiology Research Fund, Perfect Shape Medical Limited, Kai Chong Tong, Tse Kam Ming Laurence, Foo Oi Foundation Limited, Betty Hing-Chu Lee, Ping Cham So, and Lo Ying Shek Chi Wai Foundation. The funding sources had no role in the study design, data collection, analysis, interpretation, or writing of the report.

