## supplementary files for "Persistent lung inflammation and alveolar-bronchiolization due to Notch signaling dysregulation in SARS-CoV-2 infected hamster"

FigS1

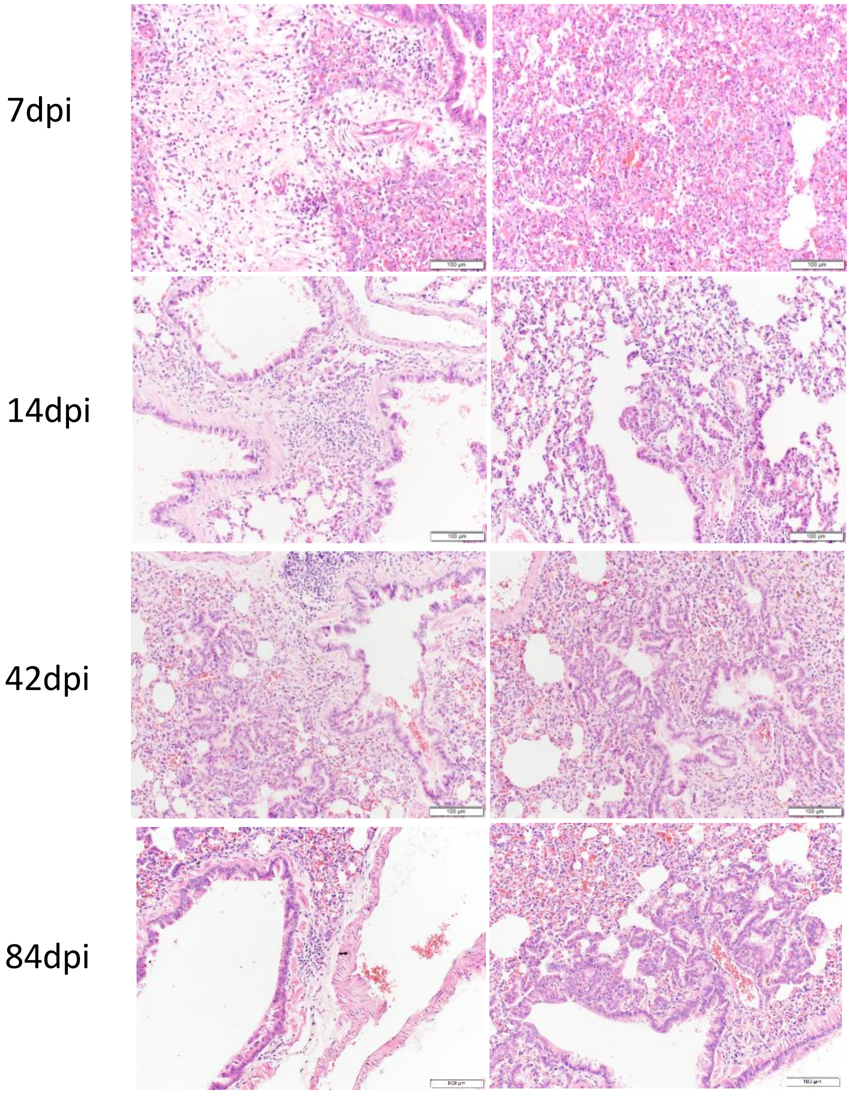


**Figure S1. SARS-CoV-2 infection causes inflammatory infiltration in hamster lungs.**
Representative H&E images showing inflammatory infiltration in SARS-CoV-2 infected hamster lungs at 7, 14, 42 and 84dpi. Scale bar=100µm.

FigS2

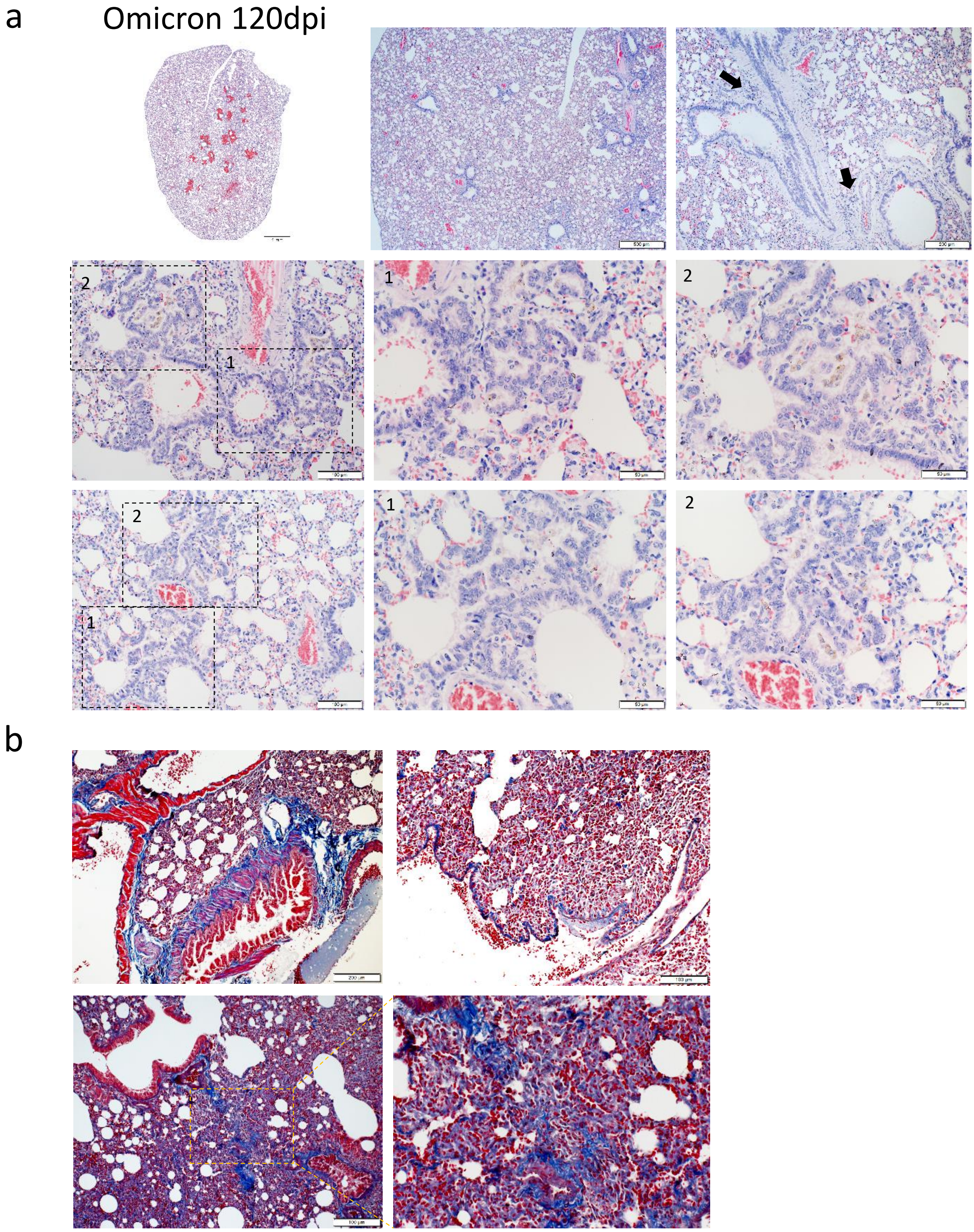


**Figure S2. Omicron BA.5 causes persistent inflammation, alveolar-bronchiolization and fibrosis in hamster lungs.**

a. Representative H&E images of Omicron BA.5 infected hamsters at 120dpi. Black arrows indicated infiltration. b. Representative images of masson trichrome stained lung of Omicron BA.5 infected hamsters at 120dpi. Scale bar=1mm, 500, 200µm, 100µm or 50µm.

FigS3

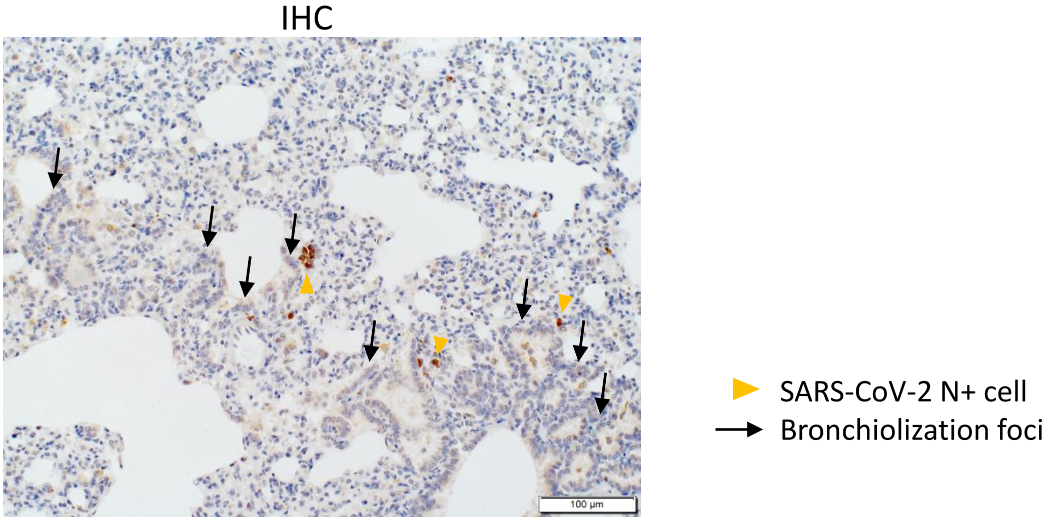


**Figure S3. Sustained virus persistence in hamster lungs after SARS-CoV-2 infection.**
Representative image of Immunohistochemistry stained SARS2-CoV-2 N protein around bronchiolization foci in SARS-CoV-2 infected hamster lung at 42dpi. Black open arrows indicated bronchiolization foci. Yellow triangles indicated SARS2-CoV-2 N protein positive cells.

FigS4

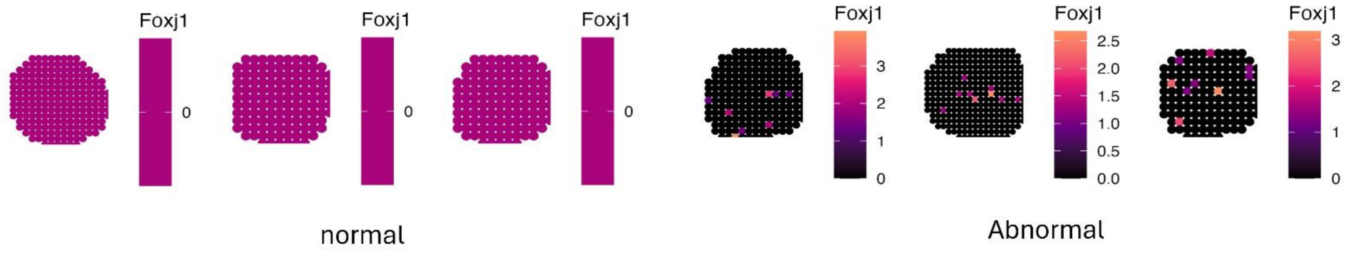


**Figure S4. Expression of gene *Foxj1* detected on abnormal bronchiolar circle regions.**

FigS5

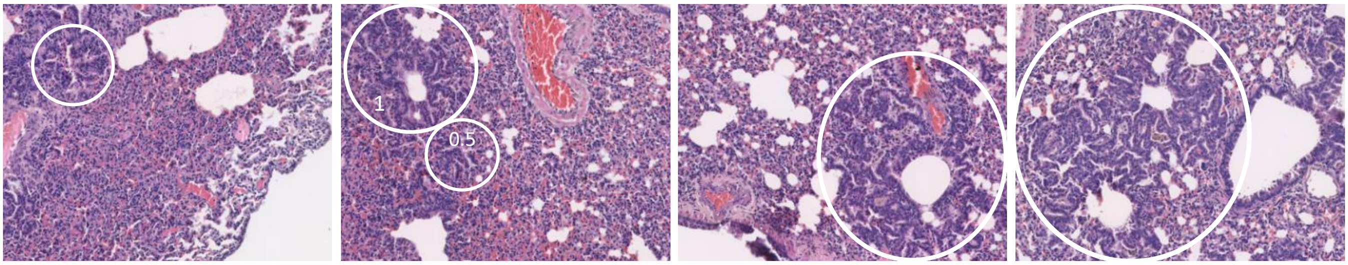


Score 0.5
Incomplete structure
or developing foci

Score 1
Less than ¼
or 1-2 foci fusion

Score 2
¼- ½
or 3-4 foci fusion

Score 3
More than ½
or more than 5 foci fusion

**Figure S5. Semi-quantification scoring of the bronchiolization foci.**
Representative image of Score 0.5, 1, 2, 3.
Tissue full slide scan was captured with Akoya Vectra Polaris Scanner. Semi-quantification was assessed using QuPath. Under each 100× view, bronchiolization foci were scored as follows: a score of 3 is assigned to foci that encompass more than 1/2 of the view or exhibit more than 5 foci fusion; a score of 2 is given to foci covering between 1/4 and 1/2 of the view or displaying 3-4 foci fusion; a score of 1 is allocated to foci occupying less than 1/4 of the view or showing 1-2 foci fusion; a score of 0.5 is assigned to incomplete structures or developing foci. The semi-quantification score is calculated using the following formulation: Total score = (0.5 * number of level 0.5) + (1 * number of level 1) + (2 * number of level 2) + (3 * number of level 3).

**Supplementary Table 1. Primer list**

| Gene name | Forward primer (5’ to 3’) | Reverse Primer (5’ to 3’) |
|---|---|---|
| *SARS-CoV-2 RdRp* | CGCATACAGTCTTRCAGGCT | GTGTGATGTTGAWATGACATGGTC |
| | Probe (5’ to 3’): FAM-TTAAGATGTGGTGCTTGCATACGTAGAC-lABkFQ | |
| *IAV M* | CTTCTAACCGAGGTCGAAACG | GGCATTTTGGACAAAKCGTCTA |
| *β-actin* | ATGGCCAGGTCATCACCATTG | CAGGAAGGAAGGCTGGAAAAG |
| *IL-6* | TGTCTTCTTGGGACTGCTGC | CCAAACCTCCGACTTGTTGA |
| *TNF-α* | CACCCACCGTCAAGGATTCA | TTGGCTGGGCAATGAAGAGT |
| *IFN-γ* | ATGGAGGGGACCTCGTCTTT | GATGGCCTGGTTGTCCTTCA |
| *CXCL10* | TACGTCGGCCTATGGCTACT | TTGGGGACTCTTGTCACTGG |
| *CCL3* | GGTCCAAGAGTACGTCGCTG | GAGTTGTGGAGGTGGCAAGG |
| *IL13* | GGTTCTCTTCCTTCGCCCTG | GCCCTCTGGTCTTGTGTGAT |
| *Fgβ* | GGTGGGGGAAAACAGAACCA | CATTGGGATTGGCTGCATGG |
| *IL33* | GCAGAAGGGGAGGCAGAAATCA | GAGATGTGGGAGGCAGGATTG |
| *FGF1* | TTCTACTTGAGAAGGCAGCAG | TCTGTTGTGGGAGCCTTTCTT |
| *FGF2* | TCTCCTGTTTCTGTCCGCTTC | AGTAATACGAAAGCTGGGGGA |
| *FGF7* | GAGTCCAGAGCAAACGGCTA | TCACCTTGCCTCGTTTGTCA |
| *MMP2* | CCATTTGATGGCAAGGATGGA | CATACTTTACCCGGACCACTTG |
| *MMP9* | TCTTCCAGTACCAAGACAAAG | AGGAAGTCGTAGGTCATGTAG |
| *Tgfβ1* | CGGGATCAGCCTCAAACG | TGAGGAGCAGGAAGGGTCTGT |
| *TIMP1* | AAAGGATTCGATGCGGTGGG | GTCCCGCGATGAGAAACTCC |
| *TIMP2* | GCCCCATGATCCCATGCTAC | TCGCTTCTCTTGATGCAGGC |
| *TIMP3* | ACTTGCCTTGCTTTGTGACC | CGGATGCAGGCATAGTGTTTG |
| *Hes1* | ACAAACCAAAGACCGCCTCT | TTGGAATGCCGGGAGCTATC |
| *Hey1* | GCCCTGGCTATGGACTATCG | TGGGAGGCGTAGTTGTTGAG |
| *Jag1* | CATCCGCGACGAGTGTGATA | CTGAAAGGCAGAACGATGCG |
| *Notch1* | ACCCTTGCTACAATCAGGGC | GCCACCTGTGAAGCTGTAGT |
| *Notch2* | TGTTAGCCCACCACCACATC | CACGGTTCATCCAGTCTGCT |
| *Notch3* | CGGGACCCCAACTACTCAAC | TCAAGTAAGGGTGCTCGCTG |
| *Sox2* | AGTGGTACGTTAGGCGCTTC | ACCCAGCAAGAACCCTTTCC |
